# Genomic rearrangements rewire chromatin contacts and drive adaptive gene expression divergence in mammals

**DOI:** 10.1101/2025.08.10.669527

**Authors:** Jui Bhattacharya, Mohan Lal, Kuljeet Singh Sandhu

## Abstract

Genomic rearrangements (GRs) are pervasive in mammalian evolution, yet their contribution to adaptive trait divergence remains poorly defined. By examining 71 mammalian genomes across eight major clades, we uncovered 206 clade-specific disruptions of ancestral synteny. These rearrangements consistently localize near genes underpinning clade-specific physiological, metabolic and behavioural traits and coincided with the accelerated expression rates in the affected lineage. The disruption of otherwise syntenic chromatin contacts and the fusion of domains with contrasting epigenetic states implied that the alteration in regulatory adjacency may underlie the expression divergence. Primate-specific rearrangements exhibited signatures of selective constraint, including fixation across species, reduced recombination, elevated linkage disequilibrium, and enhanced co-expression of genes flanking rearranged loci, indicating that these genomic architectures are selectively maintained. Our observations imply that the clade-specific genomic rearrangements repeatedly reconfigured the mammalian regulatory landscape and likely served as an evolutionary substrate for lineage-specific adaptation.

## Introduction

Genomes have gone through several rounds of structural rearrangements during the course of evolution. These rearrangements reshape the genome order and, therefore, may change the regulatory context of the genes. Such alterations may lead to expression divergence, of genes popularly known as chromosomal position effect(Wallrath and Elgin 1995; Elgin and Reuter 2013). Ectopic position effects can lead to diseases in human and mice(Milot et al. 1996; Kleinjan and Van Heyningen 1998). For example, translocation breakpoint between PAX6 and its downstream regulatory elements associates with Aniridia in human(Fantes et al. 1995). A genomic rearrangement at 100kb upstream to Steel(SI) locus downregulates Mgf gene and causes sterility in female mice(Bedell et al. 1995). Similarly breakpoints near SOX9, TWIST, and RIEG genes have association with campomelic dysplasia, Saethre-Chotzen, and Rieger syndromes respectively in human(Wirth et al. 1996; Krebs 1997; Flomen et al. 1998). Yet the evolutionary implications of chromosomal position effects remain largely under appreciated. De et al had shown that expression divergence associated with the position effects may contribute to phenotypic differences between human and chimpanzee(De et al. 2009). Bagadia et al showed that the genomic rearrangements may explain the evolutionary divergence of some brain related traits in rodents(Bagadia et al. 2019). Recently, Real et al(Real et al. 2020) identified genomic rearrangements that altered the regulatory context of gonadal gene expression governing intersexuality in moles. These studies were limited by the number of species analysed and by the context of genomic rearrangements. With the availability of large number of mammalian genomes, it is now possible to examine the phenotypic associations of genomic rearrangements in mammals. Through analysis of 71 mammalian genomes representing 8 clades, we identified 206 clade-specific long-range genomic rearrangements marking evolutionary loss of synteny. We found that the clade-specific breakpoints flanked by the genes associated with specific phenotypes of respective clades. The clade-specific breakpoints frequently disrupted the conserved enhancer-promoter communications, which coincided with the diverged expression rates of the flanking genes. Our observations implied the wide-spread adaptive significance of genomic rearrangements in mammals.

## Results

By applying the strategy outlined in the Methods section, we first identified all pairwise instances of long range (>10 Mb and *trans*) genomic disruptions of synteny across 71 mammals(Figure 1). The phenogram and the Disparity-Through-Time (DTT) plots of mean GR frequencies suggested divergence of evolutionary rate of GRs in artiodactyls, cetaceans, and rodents when compared to the rates expected under Brownian Motion (BM) model (Figure 2a-b). Following the strategy given the methods section, we further identified 206 clade-specific evolutionary breakpoint regions (EBRs) engaged in loss-of-synteny (LoS) events (Figure 2c). Rodents and artiodactyls exhibited the most numbers of clade-specific loss-of-synteny (cs-LoS) events (94 and 38 respectively), which were consistent with the earlier reports (Luo et al. 2012; Kim et al. 2017). More than 90% of gene-pairs flanking clade-specific breakpoints were syntenic in the eutherian ancestor(Kim et al. 2017), validating our inference of ancestrally syntenic configuration of the identified cases (Figure S1)

**Figure 1.**
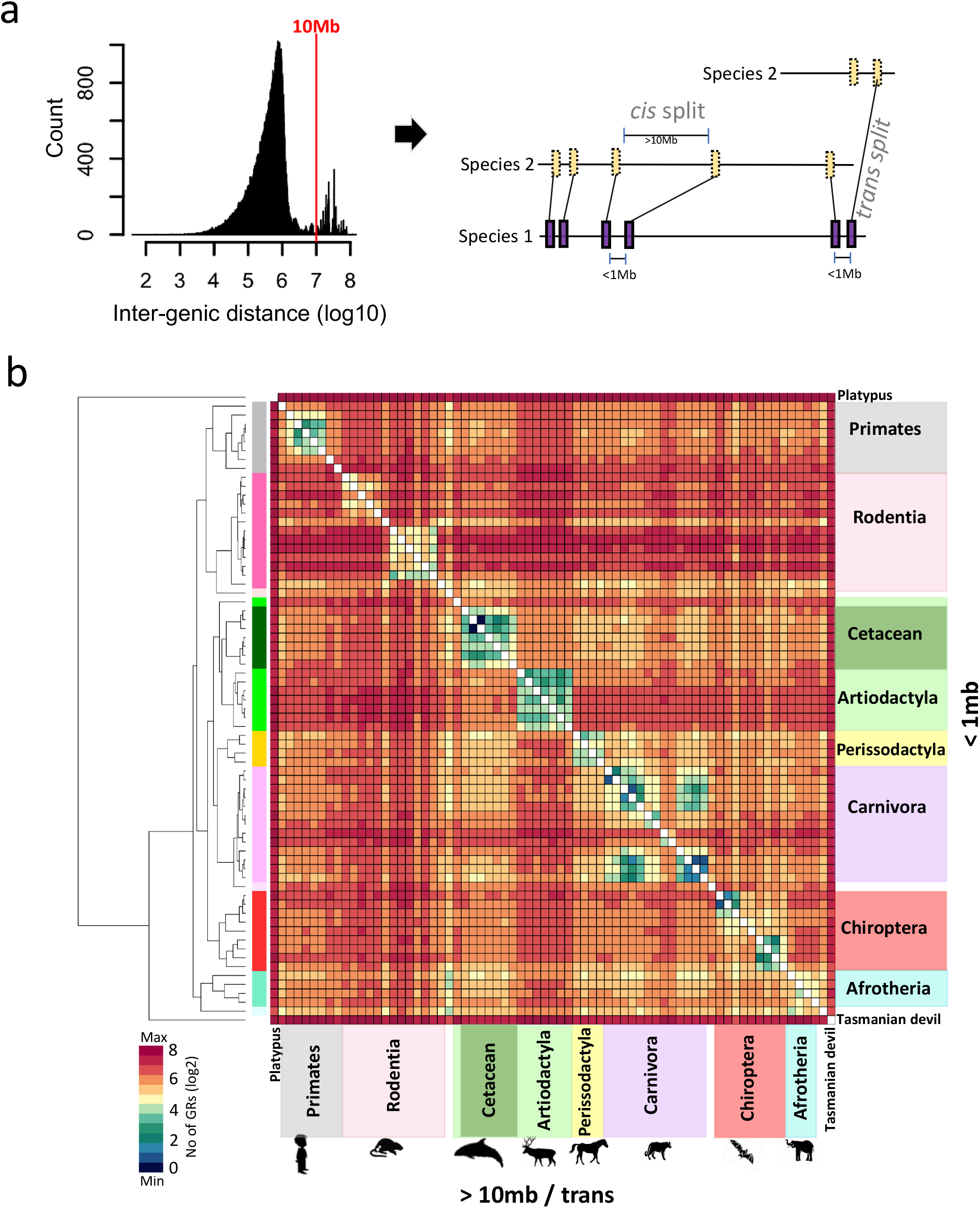
Mapping long-range genomic rearrangements across mammalian lineages. **(a)** Schematic representation of the strategy to map long-range genomic rearrangments. The distribution plot of genomic distance between adjacent genes in a species, which were <1 Mb apart in another species. Vertical line demarcates the separation of gene-pairs by >10mb in a species. **(b)** Heatmap representation of number of pair-wise long-range genomic rearrangements between lineages

**Figure 2.**
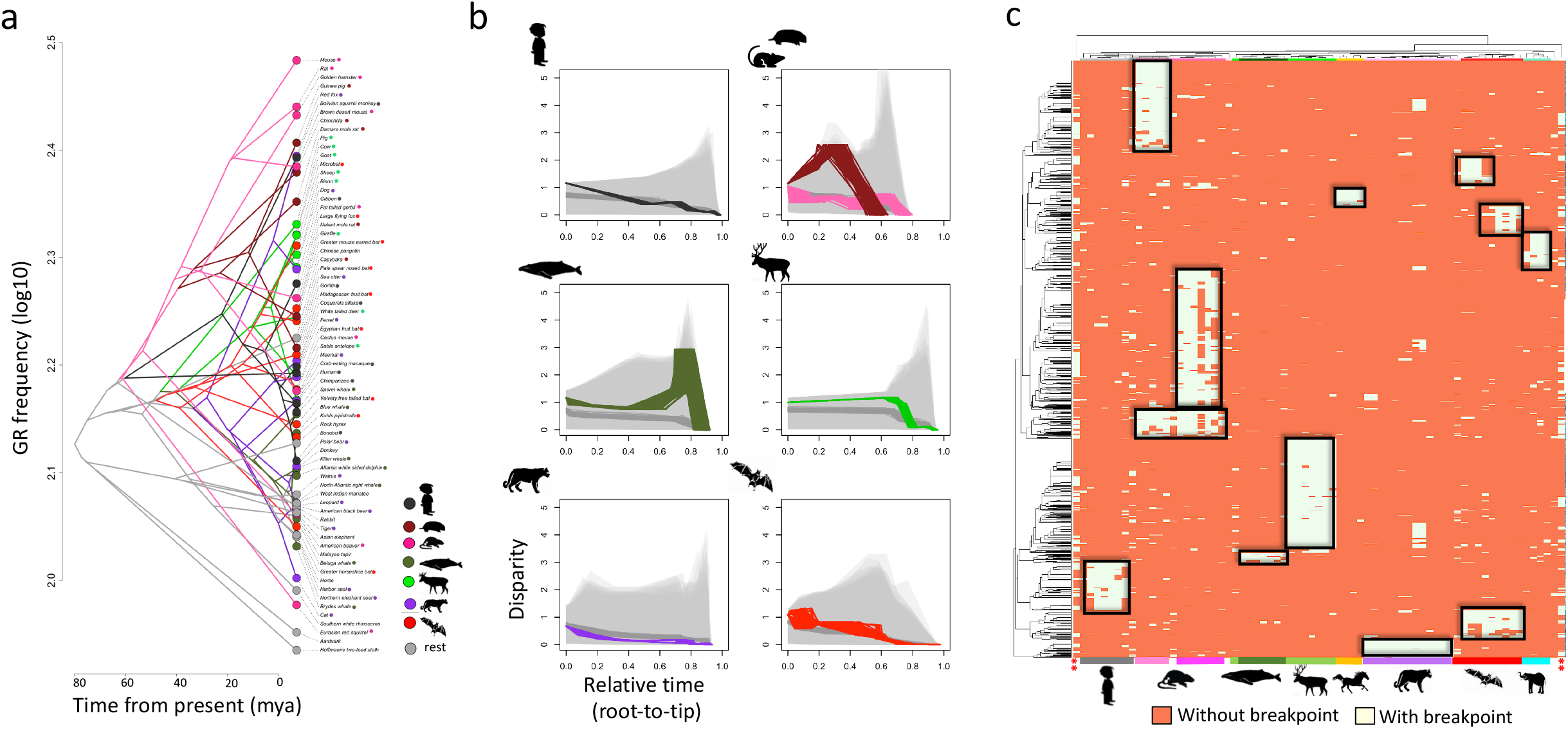
Phylogenetic analysis of GR frequency. **(a)** Phenogram of mean GR frequency across mammals. The nodes and edges are coloured clade-wise. Ancestral nodes are inferred using Maximum Likelihood approach **(b)** Clade-wise Disparity-Through-Time (DTT) plots. The observed trait variations are coloured clade-wise. The expected variations (dark grey) are simulated assuming Brownian Motion model. Light grey curves represent the 95% confidence interval of simulated variations. **(c)** Clade-specific loss of synteny (LoS) events inferred using strategy given in the Methods section. Highlighted in rectangles are the clade-specific LoS events.

To assess if the cs-LoS EBRs position preferably near certain functional loci, we obtained the genes within and the flanking 10 genes on the either sides of cs-LoS EBRs. Gene ontology analysis yielded enrichment of clade-specific functional terms. Primates exhibited enrichment of glucocorticoid secretion (CRHR1, TAC1, CRY1 etc.), lipid transport, synaptic transmission and behaviour related terms (SHANK2, RELN, FOXP1, GIT1, KLHL1, NRG1, MAPT etc.) (Figure 3). These functions have associations with primate-specific trait divergence(Bozek et al. 2015; Muntané et al. 2015; Dunn-Fletcher et al. 2018; Spanaki et al. 2024). The genes flanking artiodactyl-specific LoS break-points were enriched with flavonoid and lipid metabolism related terms, including genes like UDP-glucuronosyltransferase 1A (UGT1A) and Fatty Acid Binding Protein 4 (FABP4) (Figure 3). These terms associate with vegetarian diet and musculoskeletal health respectively, which coincide with herbivory and relatively large body mass of artiodactyls(Jiang et al. 2014; Kawai et al. 2021). Carnivore-specific LoS exhibited enrichment of extra-cellular matrix proteins, particularly matrix metalloproteases (MMPs) (Figure 3). It is established that carnivores, unlike other mammals, have endotheliochorial placenta(Wooding and Burton 2008). Differential expression of MMPs is the hallmark of invasive placentation in carnivores(Diessler et al. 2017). We further hypothesize that enrichment and expression divergence of MMPs in carnivores may also relate to effective wound healing in carnivores being susceptible to frequent wounds due to predation and the territory defence. Rodents exhibited enrichment of renal filtration related terms, aligning to their higher glomerular filtration rate, which is an adaptation to physiological requirements in diverse habitats and dietary habits of rodents(Dickinson et al. 2007; Benjamin et al. 2015; Bittner et al. 2021). The term includes the genes Anillin, Glucocorticoid Induced 1 (GLCCI1), Thrombospondin type-1 domain-containing 7A (THSD7A), Apolipoprotein L1 (APOL1) etc., which have strong associations with renal filtration and implicate in kidney diseases(Nishibori et al. 2011; Gbadegesin et al. 2014; Tomas et al. 2014; Pays 2021). Top enriched genes in Chiroptera-specific LoS break-points exhibited immunity related functions, adhering to pronounced immune tolerance in bats(Sulak et al. 2016; Jebb et al. 2020). Among others, we also observed genes like CEBPA and TNFRSF1A that are associated with echolocation in bats(Hudson et al. 2014; Wang et al. 2019). The cetacean break-points show weak enrichment of r-RNA processing and sterol related functions, aligning to longevity, body mass and diverged metabolism of cetaceans(Doherty and de Magalhães 2016; Derous et al. 2021) Afrotheria-specific LoS showed moderate enrichment of potassium ion channels, vitamin-K metabolism, and the blood coagulation (Figure 3). Ion transporters are linked to the cell proliferation, cell size and cancer, the traits significantly diverged in afrotheria clade(Savage et al. 2007). Vitamin-K is linked to the osteoblast differentiation, likely associated with the larger body mass of afrotherians like elephant, manatee etc. Blood coagulates faster in elephants(Lewis 1974) and the presence of blood coagulation factors in our set might mark an adaptation against an ancient fatal haemorrhagic disease caused by elephant endotheliotropic herpesvirus(Guntawang et al. 2021). Similarly, perissodactyl-specific breakpoints showed moderate enrichment of eye development and aerobic respiration, which lines up with the large size, 350^°^ visual field, unique motion sensitivity of horse eye(Roth and McGreevy 2025), and remarkably high aerobic respiration capacity of equine lungs(Poole and Erickson 2011) (Figure 3).

**Figure 3.**
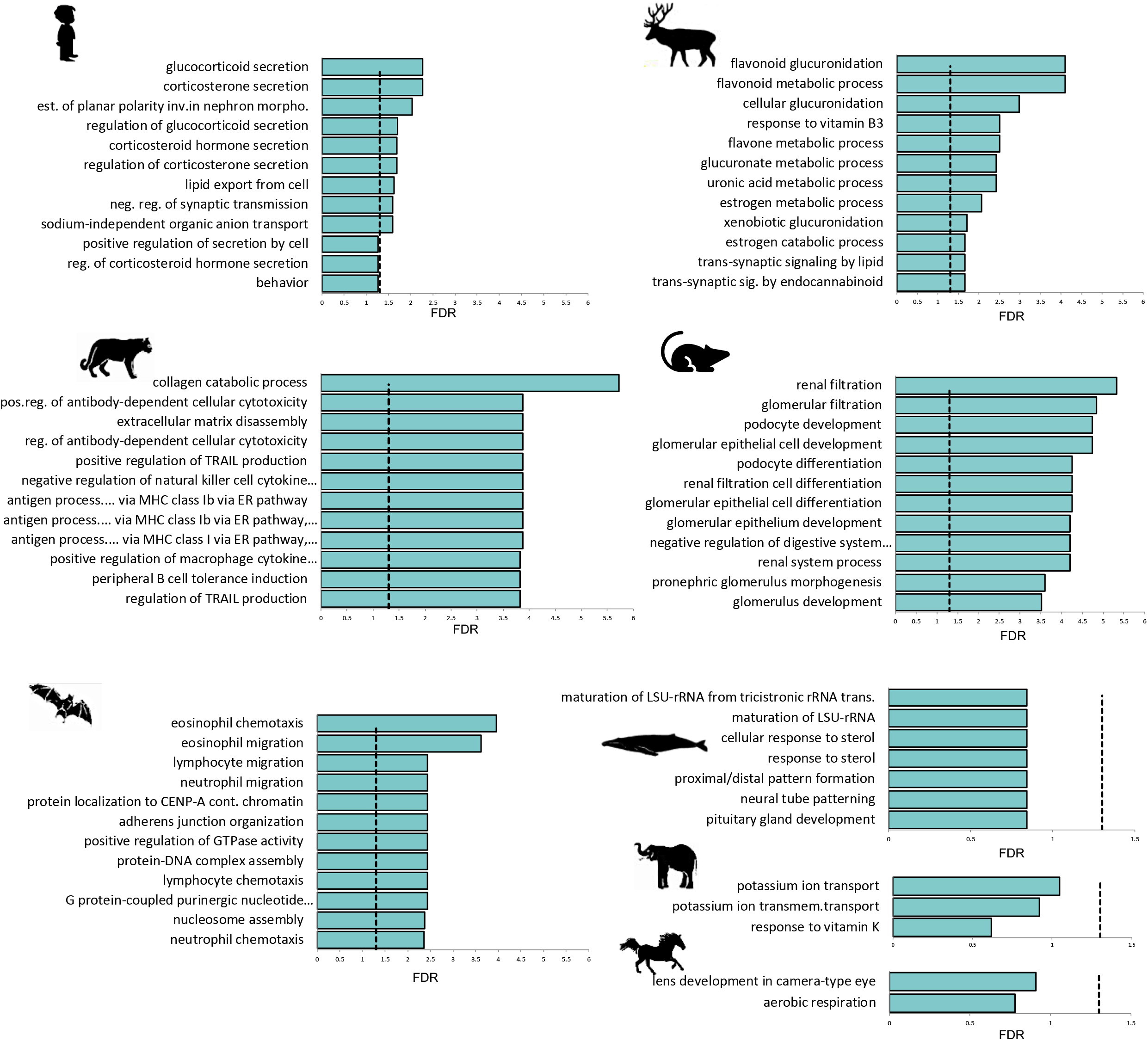
Functional associations of synteny disruptions. Enrichment of GO Biological Process terms among genes flanking the clade-specific EBRs. We implemented Benjamini-Hochberg correction of multiple testing. FDRs on the x-axis are converted to -log10 scale. The vertical lines mark the FDRs of 0.05.

In most clades, genes proximal to the cs-LoS breakpoints exhibited the expression divergence from that of the phylogenetically proximal clade controlled through an outgroup, implying the adaptive significance of genomic rearrangements (Figure 4).

**Figure 4.**
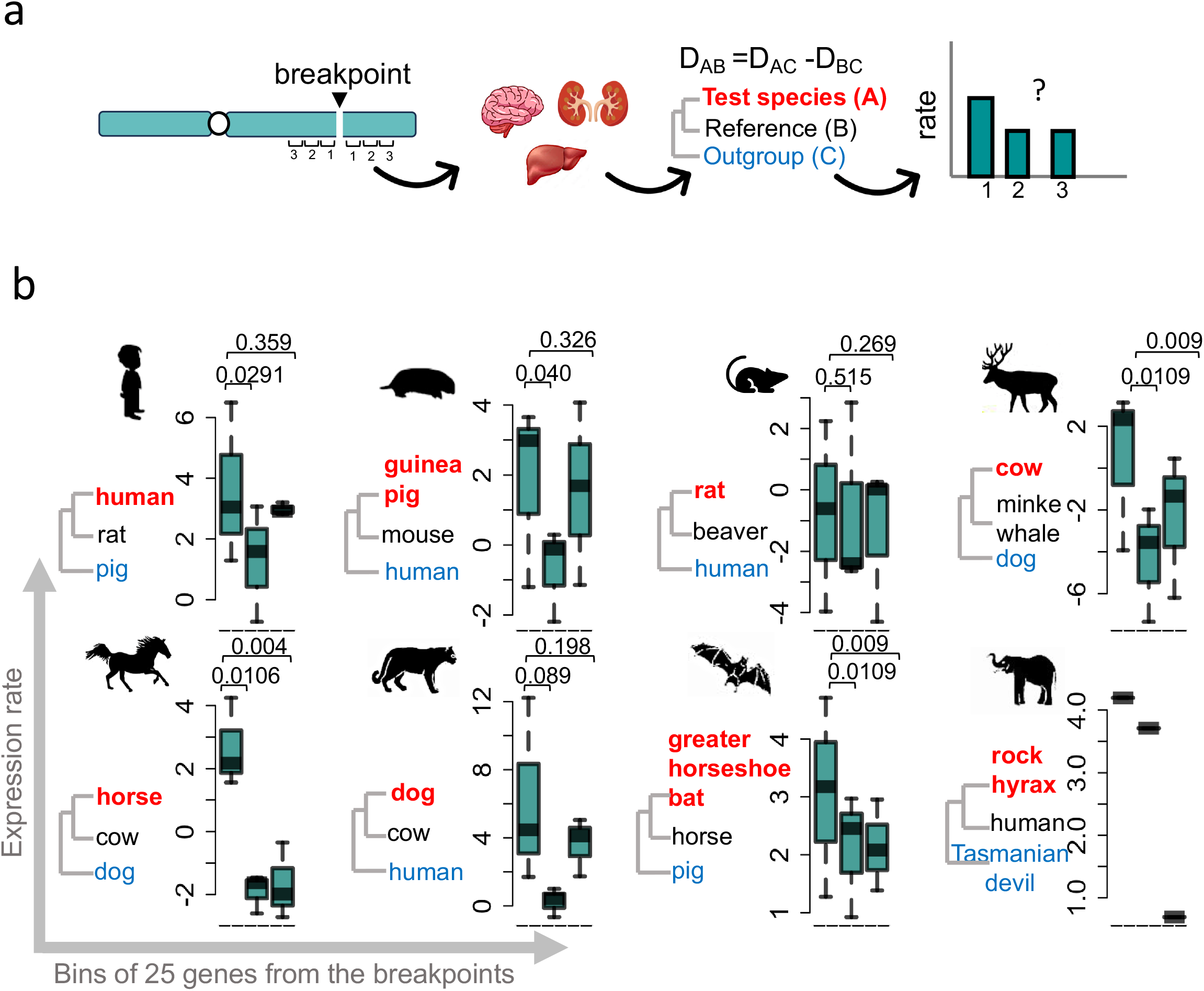
Gene expression evolution around clade-specific breakpoints. **(a)** Overall strategy of expression rate analysis. Three consecutive genomic bins as function of distance from the clade-specific breakpoints were taken. Each bin contained 25 genes. Expression rates of genes each bin was calculated assuming Ornstein-Uhlenbeck model of transcriptome evolution. **(b)** Lineage-specific expression rates of the genes as a function of distance from the clade-specific evolutionary break-points. Each box represents the distribution of expression divergence values of 25 genes incrementally distant from the breakpoint. A species from the nearest order or sub-order was taken as reference species, while a species hierarchically outside the clade of test and the reference species was taken as outgroup species. P-values were calculated using Mann-Whitney U tests.

The chromosomal position-effect may explain the observed expression divergence. To test this, we first analysed the chromosomal coordinates of enhancer-promoter pairs that exhibited evolutionarily conserved synteny. We observed non-random accumulation of clade-specific breakpoints within enhancer-promoter spans, which were syntenic in other clades, suggesting that the expression divergence of genes might have been driven by the loss of proximity to their cognate enhancers (Figure 5a-b). Secondly, we assessed the position effect through analysis of chromatin compartments. We observed that the chromatin compartments with contrasting states (A and B) in a species tended to fuse together and subsequently exhibited a coherent state (A or B) in the species exhibiting cs-LoS event (Figure 5c-d). This is marked by a decrease in Kronecker’s delta (Methods) of consecutive genomic bins around breakpoints (Figure 5c). This may signify an alteration in gene expression through the mixing of contrasting epigenetic states *via* gained proximity. We illustrated this through an example of primate-specific genomic rearrangement near Neurokinin-1 (TAC1), a glucocorticoid secretion and behaviour related gene. The gene locates in the B-compartment in mouse, which gains proximity to an A-compartment, and subsequently switches from B-to-A compartment in neuronal cells (Figure 5d). Overall the split of enhancer-promoter synteny and gain of proximity between A and B chromatin compartments highlight the mechanistic basis of expression divergence of genes located near cs-LoS breakpoints.

**Figure 5.**
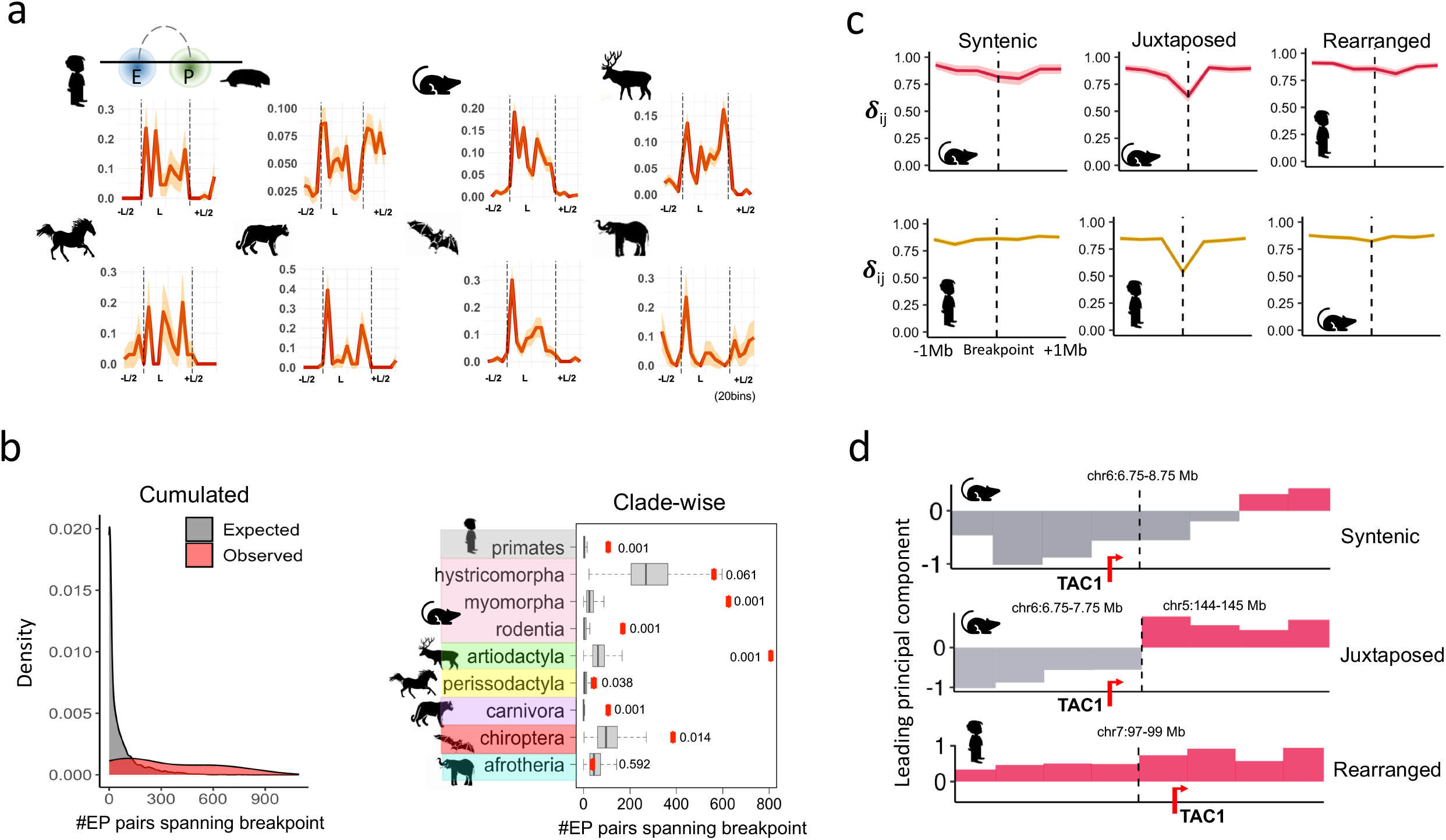
Chromosomal position effect associated with the clade-specific loss of synteny. **(a)** Enrichment of clade-specific breakpoints within and around enhancer-promoter loops in the species having syntenic configuration. Loop coordinates were extended either side by half of the loop length-L. **(b)** The number of conserved enhancer-promoter contacts that had lost proximity through GR in a clade-specific manner, superimposed onto the random null distribution obtained from controlled randomized of breakpoints. P-values are calculated using bootstrap method. Shown are the plots for the cumulated set and the clade-specific sets. **(c)** Kronecker’s δ_ij_, signifying change in A and B compartment states of neighbouring domains, as a function of distance from the break-points in primates and rodents. Drop in Kronecker’s delta signifies the switch in states (A-to-B or B-to-A). The three plots correspond to syntenic configuration in mouse (left), distal regions (syntenic in human) when juxtaposed in mouse (middle), rearranged syntenic regions in human (right) respectively in case of primate-specific LoS. Similarly, for rodent-specific LoS, syntenic configuration in human (left), distal juxtaposed regions in human (middle) and rearranged syntenic regions in mouse (right) are plotted. **(d)** Chromatin compartment states (leading Eigen vector of Hi-C contact matrix) for regions around TAC1 locus, which exhibited primate-specific LoS nearby, in mouse and human neuronal cells. A and B compartments are coloured as red and grey colours respectively.

We further exemplified a primate-specific LoS event near Corticotropin Releasing Hormone Receptor 1 (CRHR1) gene that coordinates the autonomic, endocrine and behavioural response to stress and depression through Hypothalamus-Pituitary-Adrenal (HPA) axis(Jezova et al. 1999) and may also regulate the timing of birth(Smith and Nicholson 2007). Interestingly, CRHR1 shows primate-specific conservation of 29 amino acids of exon-6, and insertions of retroviral LINE elements at the upstream region (Figure S2). The gene also exhibits loss of chromatin contacts and expression divergence in primates (Figure 6a-b). We propose that the sequence and the expression divergence of this locus may link to primates’ adaptation to stress from complex social structures(Pusey et al. 1997; Sapolsky 2021). Similarly, artiodactyl-specific LoS near UGT1A genes, which metabolizes flavonoids commonly present in vegetarian diet, exhibited disruption of chromatin contacts and expression divergence in artiodactyls (Figure 6c-d). UGT1A genes have expanded copy numbers in herbivores(Kawai et al. 2021), implying convergence of distinct molecular mechanisms underlying expression evolution of UGT1A genes in artiodactyls.

**Figure 6.**
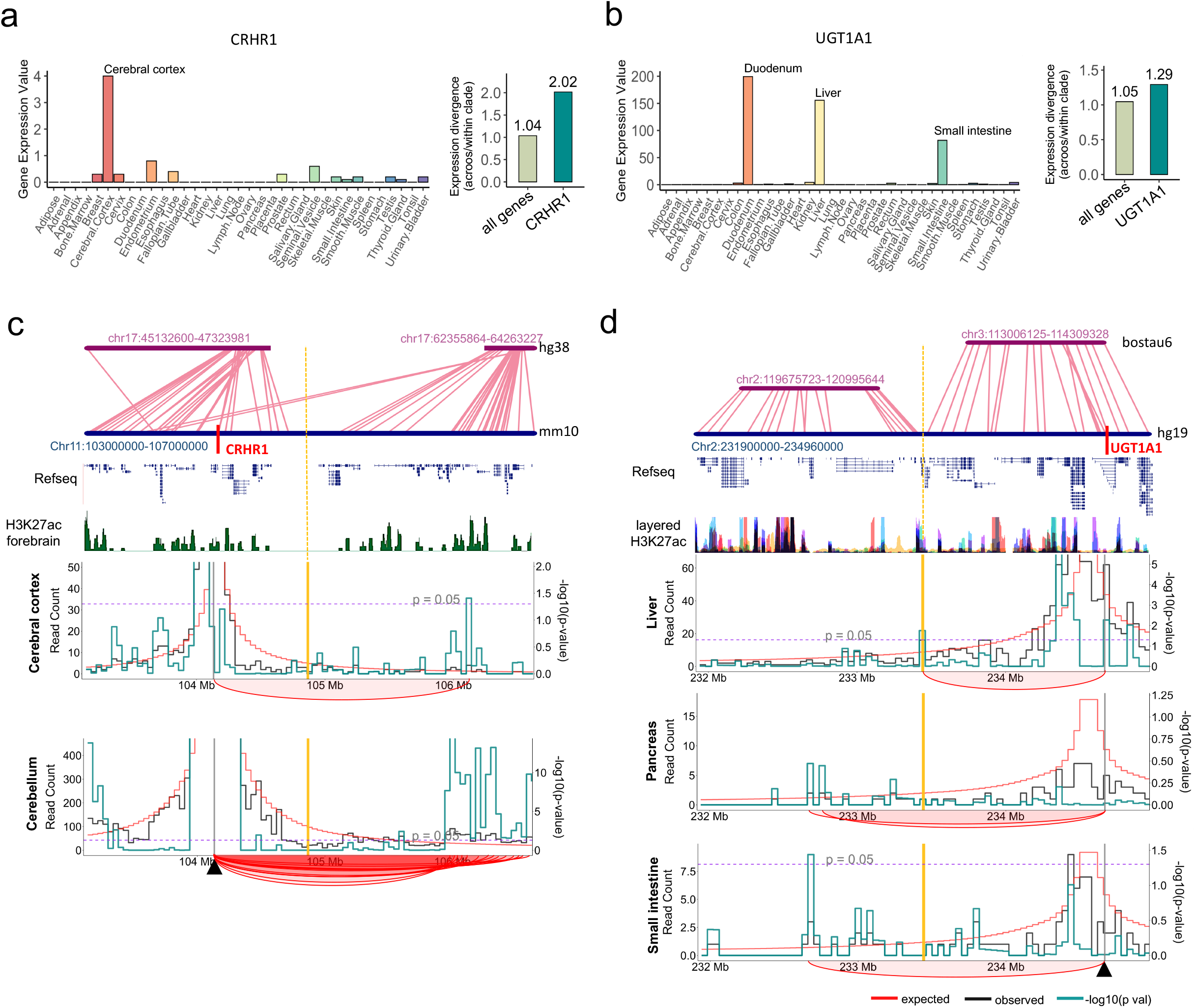
Illustrative examples of clade-specific breakpoints. **(a-b)** Left-panels: CRHR1 and UGT1A expression values in 35 human tissues respectively. Right-panels: expression divergence calculated as the Euclidean distance across tissues and summarized as ratio of across-clade to within-clade distances. **(b-c)** Mouse-to-human and human-to-cow synteny plots around (b) CRHR1, and (c) UGT1A gene respectively. Shown below are the tracks of RefSeq non-redundant genes, H3K27ac modification, and virtual 4C in relevant tissues. The clade-specific break-point regions are highlighted by vertical bars. 4C reference loci are highlighted using arrow-heads. Black and red curves represent observed and expected 4C signals respectively. Green curve represents -log10 of p-values. Blue dashed line represents the p-value cut-off of 0.05. Red arcs mark the significantly frequent chromatin contacts.

To assess whether the clade-specific GRs are shaped by selection, we focused on primate-specific LoS events, leveraging the available population genomic data. We found that rearranged loci exhibit elevated linkage disequilibrium and reduced recombination rates around primate-specific fusion points in humans, indicating tight genetic linkage (Figure 7a-b). These rearrangements also displayed fixed genomic adjacencies across primates (Figure 7c). In addition, genes flanking primate-specific fusion points show stronger co-expression compared with genes near corresponding primate-specific LoS breakpoints in a non-primate species (mouse) (Figure 7d). Together, these patterns are consistent with the selective constraints acting on primate-specific GRs and may extend to other clades.

**Figure 7.**
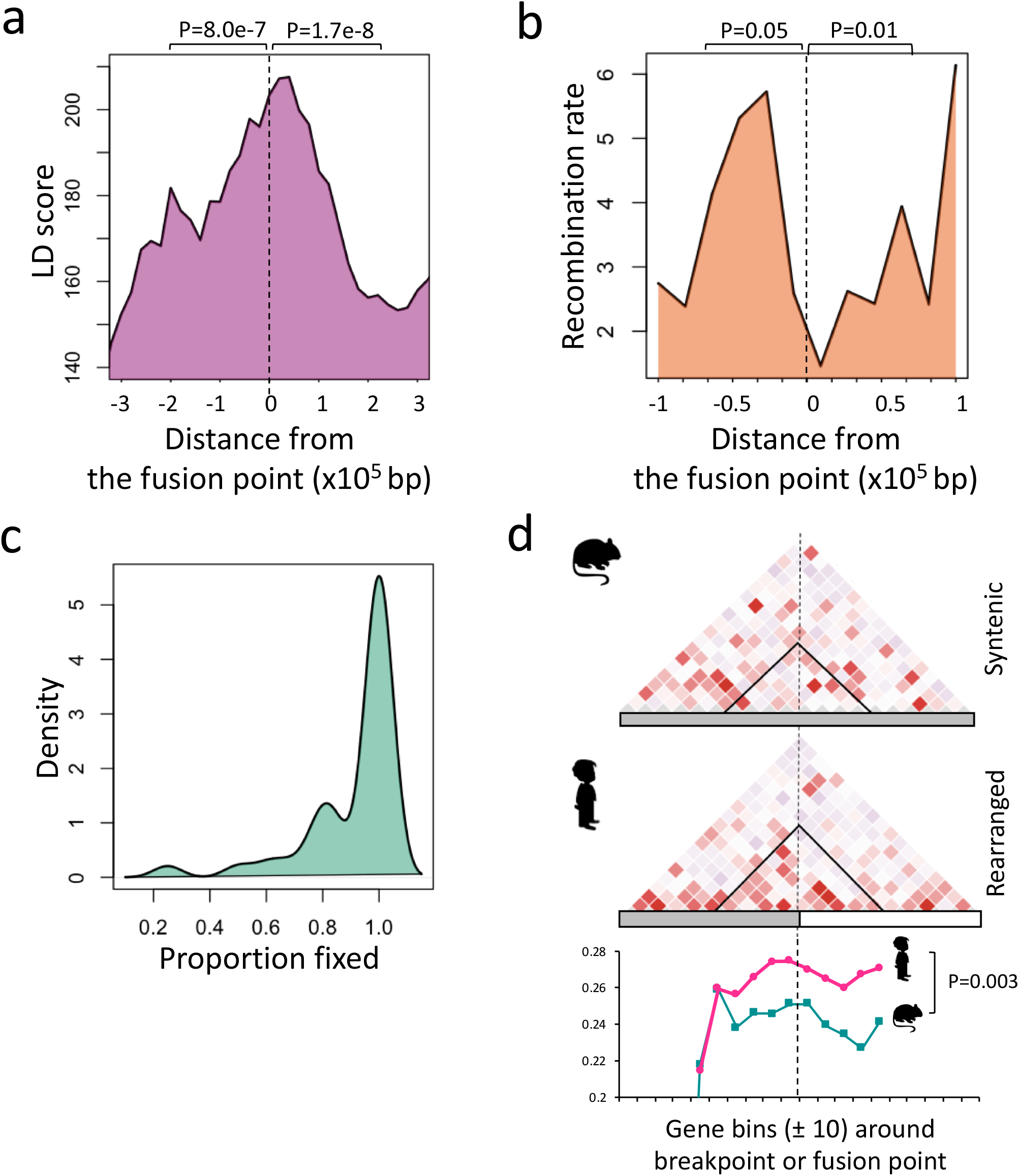
Signatures of selective constraints. **(a)** Human LD score and **(b)** recombination rates around the fusion points resulting from primate-specific LoS events. **(c)** Distribution of number of primates exhibiting evolutionarily fixed configuration of rearranged genomic regions. **(d)** Mean co-expression of genes around primate-specific breakpoints in mouse (top panel) and around rearranged fusion points in human (middle panel). Bottom panel shows sliding means of the values around diagonal of the co-expression matrices of mouse and human. P-values were calculated using two-tailed unpaired t-tests.

## Discussion

Genes are not autonomous transcriptional units within the genomic landscape but rather operate in a highly context-dependent manner influenced by their local and global chromatin environment. Reporter integration in proximity to enhancer strengthens the expression, while that within lamina-associated domains attenuates the transcriptional activity(Akhtar et al. 2013). Distinct linear and spatial positions of a gene also confer variation to its expression noise(Chen and Zhang 2016; Singh et al. 2016). The alterations in genomic context through chromosomal rearrangements can, therefore, influence the epigenetic and transcriptional states of the genes. Although studied in somatic human cells and cancer tissues(Chiang et al. 2017; Xu et al. 2022), the evolutionary dynamics of such position effects remains poorly understood. Our observations through this study highlight that the clade-specific long-range genomic rearrangements associate with the enrichment and expression-divergence of genes with clade-defining traits. Interestingly, these genes have also lost the proximity to their cognate enhancers as reflected from the splitting of enhancer-promoter pairs, likely representing one of the possible causes of the observed divergence of gene expression in the respective clades. The gained proximity of chromatin domains with opposing epigenetic states further strengthens the notion of chromosomal position effect in evolutionary context. We propose that the alteration in regulatory context may have contributed to the expression evolution of genes proximal to clade-specific EBRs. From our observations, it follows that clade-specific GRs may have guided the evolution of clade-specific mammalian phenotypes through gene expression evolution. Primate-specific fixation and tight linkage of rearranged loci further implied the signatures of selective constraints.

The reported genomic rearrangements and the engaged loci also offer a valuable catalogue for biomedical interpretation. Many of the reported loci also implicate in human diseases and understanding the rewiring of their regulatory contexts could help in genetic prioritization and devising medical strategies to intervene the mis-regulation. Certain mammals are resistant to certain diseases, like elephant is resistant to cancer and bats are immune tolerant to viruses. Our observations may also help studying the regulatory contexts which are instrumental in disease resistance and exploit that knowledge to understand disease susceptibilities in humans.

## Conclusion

Altogether, our observations suggest that clade-specific genomic rearrangements are not merely the passive signatures of evolutionary history and speciation but are active agents reshaping mammalian regulatory landscapes. By rewiring enhancer-promoter contacts and shuffling chromatin compartments, these rearrangements generate lineage-specific expression programs that coincide with the emergence of clade-defining phenotypes. The recurrence of such events across distant mammalian clades highlights genome topology as a pervasive, yet underappreciated, substrate of adaptive evolution. Together, our results establish long-range genomic rearrangements as an important evolutionary force that has repeatedly sculpted mammalian diversity.

## Methods

### Genome data

We obtained the chromosomal coordinates of human orthologous genes for 18 mammalian species from Ensembl (http://www.ensembl.org). Fifty three genomes not represented in Ensemble were obtained from DNA Zoo (https://www.dnazoo.org/assemblies) and Bat1K Project (https://bat1k.com/). We aligned the genome FASTA files to the human genome using LASTAL(https://github.com/lpryszcz/last). Resulting maf files were processed with *‘last-split’* and converted to chain files using *‘maf-convert’* utility (https://github.com/lpryszcz/last). To obtain orthologous genomic coordinates, we used the UCSC LiftOver tool (https://genome.ucsc.edu/cgi-bin/hgLiftOver). The phylogenetic tree of the species is given in the Figure S3. The genome assembly information is provided in Dataset S1.

### Identification of clade-specific loss-of-synteny (LoS) events

To define evolutionary breakpoints of long-range genomic rearrangements, we used a distance-based approach. For each species, we first calculated linear distances between non-redundant all possible *cis* gene pairs. Gene pairs separated by ≤1 Mb were considered syntenic in a given species. The corresponding intergenic distances between orthologues of these syntenic pairs were then obtained for all other species and their distribution was visualized as histograms using the ‘hist’, function in R. The analysis revealed a bimodal distribution with two modes separated at around 10 Mb distance. This 10 Mb threshold was, therefore, used as a threshold to define loss of synteny events. Gene pairs that were within 1 mb distance in one species but exceeded 10Mb in other species were categorized as *cis*-rearrangement. If the orthologues of syntenic pairs were located on different chromosomes in another species, they were classified as *trans*-rearrangements. To minimize noise that might result from retro-transposition or assembly errors, we further obtained the set where the orthologous disruption occurred with at least 3 genes on either side of the break-point. We further calculated the inter-breakpoint distances defined as the linear distances between gene pairs forming breakpoints across all species. The distribution of distances suggested a distinct natural cut-off of 1.4 Mb (Figure S4), below which the breakpoints can be considered as clustered into evolutionary break-point region (EBRs). An EBR was identified as a clade-specific loss-of-synteny (LoS) event if it engages in LoS events in at least 50% of the species within a given mammalian clade, while being syntenic in most other clades (split in not more than two species).

### Phylogenetic comparative analyses of frequency of GRs

We used mean frequency of GRs for each species and perform ‘phenogram’ analysis using ‘phytools’ and DTT analysis using ‘geiger’ R-packges. We assumed Brownian Motion (BM) model to simulate (*nsim=1000*) the evolution of GR frequencies through time. The default average squared Euclidean distance was used as metric.

### Gene Ontology analysis

For each evolutionary breakpoint region (EBR), we extracted all genes within EBR and 10 genes upstream and downstream of both ends. These unique gene sets were compiled for each clade. Gene ontology (GO) enrichment analysis was performed using the ToppFun function from the ToppGene Suite (https://toppgene.cchmc.org/) to fetch the GO Biological Process categories. P-values were corrected using Benjamini-Hoschberg method.

### Gene expression analysis

We retrieved the expression data of CRHR1 and UGT1A1 of 35 human tissues from the TissueEnrich database(https://tissueenrich.gdcb.iastate.edu/) for tissue specific gene enrichment analysis. To calculate expression divergence, we used tissue-wide expression data from the Bgee database(https://www.bgee.org/). For each gene (CRHR1 and UGT1A1), a divergence ratio was calculated as the median of across-clade versus within-clade distances. These gene-specific ratios were compared to the genome-wide median divergence ratios calculated across all genes.

### Expression rate analysis

We used ‘TreeExp’ R-package(https://github.com/hr1912/TreeExp) to evaluate the evolutionary rates of expression divergence assuming Ornstein-Uhlenbeck model of gene expression evolution. For each clade, we assigned a test species, a reference species and an out-group species from phylogenetically adjacent clade. We made three distance-based bins, each comprised of 25 genes, on either side of the break-point. Gene expression data were obtained from Liu et al(Liu et al. 2023), which provides expression profiles for three tissues: brain, kidney, and liver for multiple species. For co-expression analysis, 10 genes on either side of breakpoint or the fusion point were obtained. Pearson’s correlation coefficients across 26 mouse tissues (https://www.ncbi.nlm.nih.gov/sra/?term=srp020526) and 55 human tissues (https://gtexportal.org/home/) calculated for gene-sets around each reference point. Mean matrix of individual correlation matrices were plotted for visualization. Means of sliding windows of 3 genes along the diagonal of the correlation matrix was used for quantitative analysis.

### Enhancer-promoter contact analysis

All human enhancer-promoter (EP) pairs and their orthologues in other species, were obtained from the dataset provided by Laverré et al(Laverré et al. 2022). The evolutionarily conserved EP pairs were then used to analyze disruption by clade-specific LoS events. If a conserved EP pair flanked a breakpoint unique to a clade, the pair was considered split by the rearrangement. For clades such as Chiroptera, Perissodactyla, Afrotheria, and the suborder Hystricomorpha, for which the enhancer orthologs were not included in the dataset by Laverré et al, the coordinates were lifted over using the UCSC liftOver tool (https://genome-store.ucsc.edu/). To calculate the statistical significance of observed EP pair disruptions, we performed bootstrap by randomizations breakpoint coordinates across the human genome 1000 times while keeping the chromosomal distribution of breakpoints same as original set. The bootstrap-based p-values were obtained using:

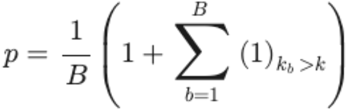

where B is the total number of bootstrap samples, k is the observed number of disrupted EP pairs.

### Hi-C analysis

We used HUGIn web browser (https://yunliweb.its.unc.edu/hugin/tutorial_page.php), to fetch virtual 4C interaction profiles at 40 kb resolution from human Hi-C data across relevant tissues. For mouse (mm10), .cool files were obtained from the 4D Nucleome Data Portal (https://data.4dnucleome.org/) and virtual 4C profiles were generated using Fit-Hi-C (https://github.com/ay-lab/fithic). We downloaded chromatin compartment (A/B) files for both human and mouse tissues from the 4D Nucleome. Change in compartment along the genomic regions spanning the break-points was calculated using Kronecker’s delta function:

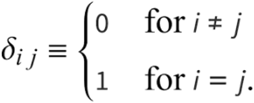

If consecutive genomic bins *i* and *j* had same compartment (A-A or B-B), the function takes the value 1, and zero when *i* and *j* bins had different compartment values (A-B or B-A). We calculated the average and 95% confidence intervals of Kronecker’s delta across all breakpoints for each tissue and clade (supplementary). For compact representation a global average of Kronecker’s delta was plotted at 250 kb resolution around +/-1 Mb to the break-point.

### Linkage disequilibrium and recombination rates

LD scores from 1000 genomes and the recombination rates were obtained from Alkes group (https://zenodo.org/records/8182036) and Halldorsson et al(Halldorsson et al. 2019) respectively. The sliding means of these scores in 20kb bins around the reference points were plotted as a function of distance from the fusion-points.

The source of all datasets used are listed in Dataset S2.

## Data availability

All data generated or analysed during this study are included in its supplementary dataset files.

## Acknowledgements

Authors duly acknowledge the financial support from Department of Biotechnology, India (BT/PR40149/BTIS/137/36/2022, BT/PR40198/BTIS/137/56/2023). JB performed all the data analysis and prepared the figures. ML helped in coding. KSS conceived, improvised, guided, and wrote the manuscript.

---

**Figure S1.**
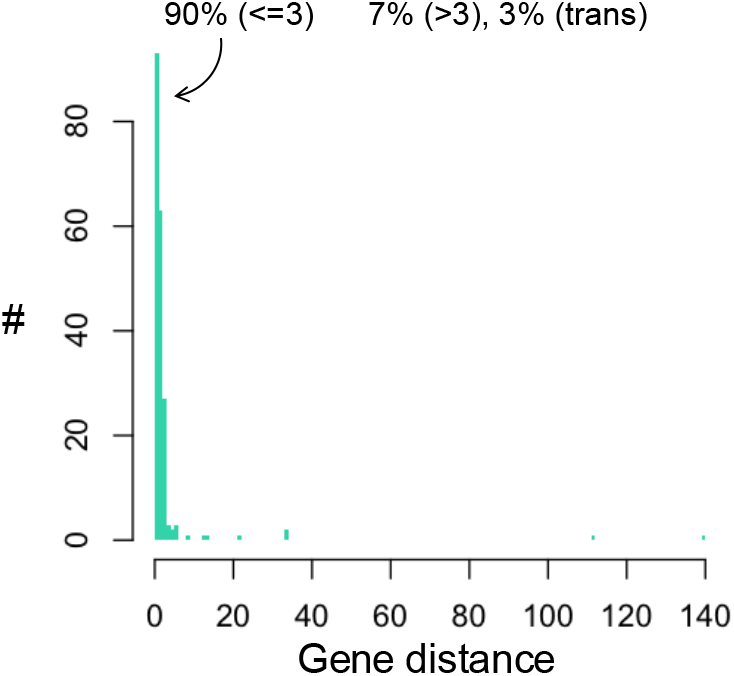
Distribution of gene-distances (#genes) between genes flanking the clade-specific breakpoints in the common ancestor of placental mammals

**Figure S2.**
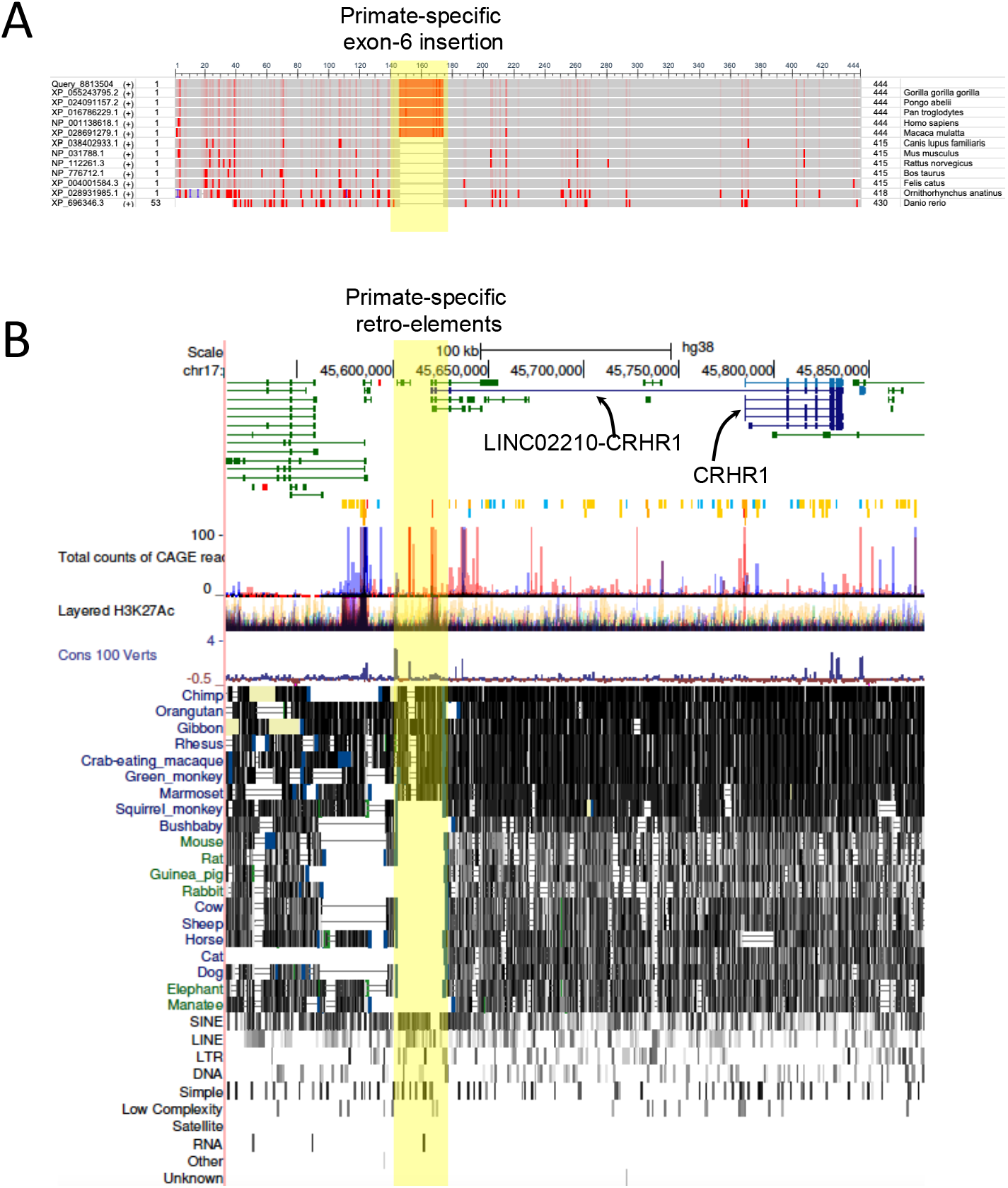
(A) Primate specific insertion of exon-6 in the CRHR1 protein. (B) Primate-specific insertion of re tro-viral elements upstream to the CRHR1 gene and at the promoter of a LINC RNA.

**Figure S3.**
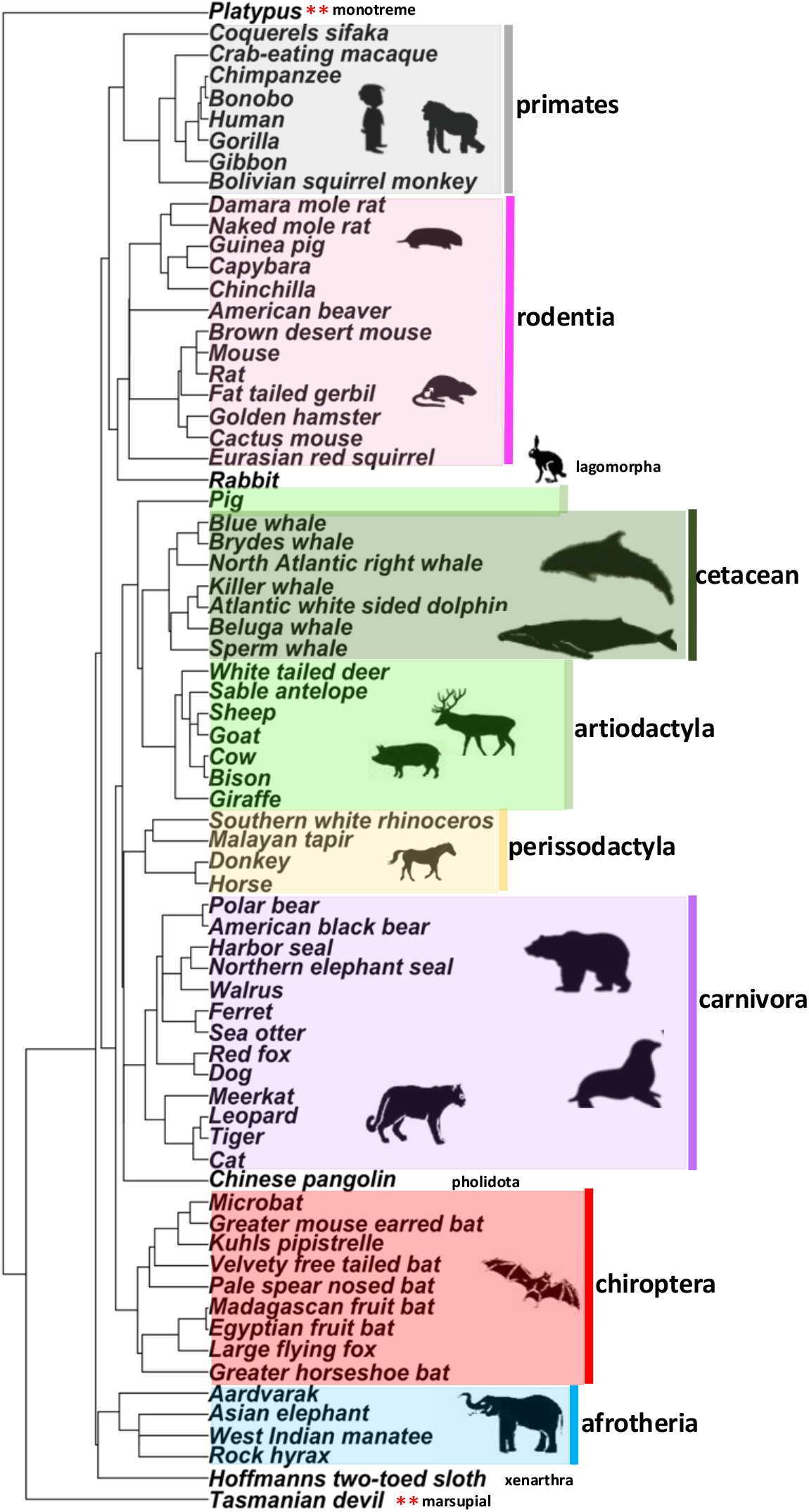
Phylogenetic tree of species analyzed in the study

**Figure S4.**
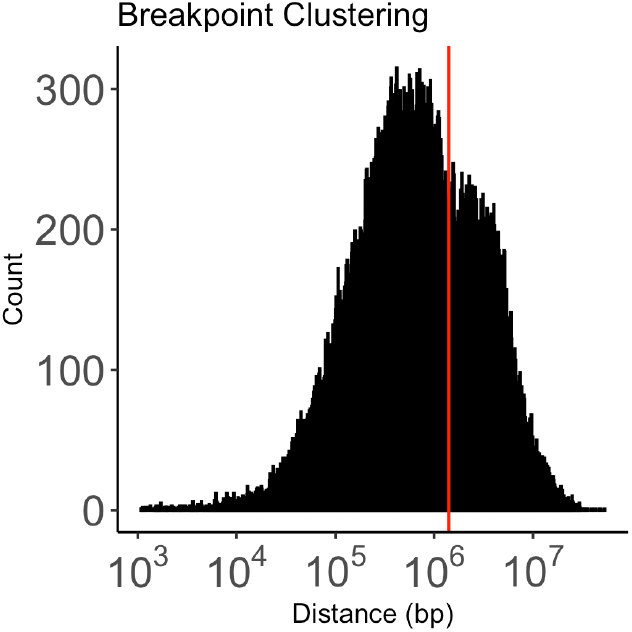
Distribution of inter-breakpoint distances across species. Vertical bar marks the cut-off of 1.4Mb

